## Supplemental Information for "Structural basis for broad anti-phage immunity by DISARM"

**Supplementary Table 1 l Cryo-EM data collection, refinement and validation statistics.**

| | **DrmAB-ADP** (PDB 7S9V, EMD-24938) | **DrmAB-ADP-DNA** (PDB 7S9W, EMD-24939) | **DrmAB** |
|---|---|---|---|
| **Data collection and processing** | | | |
| Magnification | 22,500 | | |
| Voltage (kV) | 300 kV | | |
| Electron exposure (e–/Å$^2$) | 80 | | |
| Defocus range (μm) | -1.5 to -2.5 | | |
| Pixel size (Å) | 1.1 | | |
| Symmetry imposed | C1 | | |
| Initial particle images (no.) | 4,669,932 | | 1,540,419 |
| Final particle images (no.) | 120,119 | 121,764 | 148,062 |
| Map resolution (Å) at 0.143 FSC threshold | 3.31 | 3.41 | 3.84 |
| Map resolution range (Å) | 3.1 to 4.3 | 3.3 to 4.5 | 3.6-6.8 |
| **Refinement** | | | |
| Initial model used (PDB code) | *De novo* | | |
| Model resolution (Å) | 3.6 | 3.8 | |
| FSC threshold | 0.5 | | |
| Map sharpening *B* factor (Å$^2$) | -136.8 | -151.1 | |
| Model composition<br>Non-hydrogen atoms<br>Protein residues<br>Nucleotides<br>Ligands | <br>27680<br>1808<br>0<br>1 | <br>28125<br>1751<br>7<br>1 | |
| Mean *B* factors (Å$^2$)<br>Protein<br>Nucleotides | <br>62.08<br>N/A | <br>106.66<br>116.37 | |
| R.m.s. deviations<br>Bond lengths (Å)<br>Bond angles (°) | <br>0.002<br>0.617 | <br>0.002<br>0.631 | |
| Validation<br>MolProbity score<br>Clashscore<br>Poor rotamers (%) | <br>1.52<br>3.27<br>0 | <br>1.53<br>3.34<br>0 | |
| Ramachandran plot<br>Favored (%)<br>Allowed (%)<br>Disallowed (%) | <br>93.93<br>6.01<br>0.6 | <br>93.90<br>6.1<br>0 | |

**Supplementary Table 2 | DNA oligos used for *in vitro* assays in this study.**

| Oligo name | Sequence (5'-3') |
|---|---|
| DNA stem loop | TTTTTTTCATCGATGAGCACTGCTATTCCCTAGCAGTGCTCATC GATGATTTTCATCGATGAGCGGTTTT |
| Methylated DNA stem loop* | TTTTTTTCAT**mC**GATGAGCACTGCTATTCCCTAGCAGTGCTCAT **mC**GATGATTTTCAT**mC**GATGAGCGGTTTT |
| Fluorescent DNA | Cy5-ATTTTGACAGCCCACATGGCTTGATGAGTGGCGCACTCGC CAGCCTGAGCATGGCGAAAACTCCTCCAGTCTGCT |
| Unlabeled complement DNA | AGCAGACTGGAGGAGTTTTCGCCATGCTCAGGCTGGCGAGTG CGCCACTCATCAAGCCATGTGGGCTGTCAAAAT |
| 5' block | GCCATGTGGGCTGTCAAAAT |
| 3' block | AGCAGACTGGAGGAGTTTTC |

mC corresponds to 5-methyl-Cytosine.

**Supplementary Table 3 I DNA oligos used for cloning in this study.**

| Oligo name | Sequence (5' - 3') | Used for: (in fw/rv couples) | fw/rv |
|---|---|---|---|
| BN1750 | TACTTCCAATCCAATGCAATGATCAT CAATAACAAAACTCCAG | Clone *drmB* into plasmid 13SS using LIC cloning | fw |
| BN1751 | TTATCCACTTCCAATGTTATTATTAAA AACCCTTAAAAAATGCAGC | Clone *drmB* into plasmid 13SS using LIC cloning | rv |
| BN1060 | TATATATACATATGACTGATAACAACA AATCTAG | Clone *drmA* into plasmid pACYC-Duet using RE cloning-NdeI | fw |
| BN1061 | ATATATATCTCGAGTCAATCCTCGTC TTTCGTTG | Clone *drmA* into plasmid pACYC-Duet using RE cloning-XhoI | rv |
| BN172 | AGATCTGCCATATGTATATCTCCTTC | Amplify pACYC-Duet backbone for cloning of *drmA* insert | rv |
| BN2290 | GATCGCTGACGTCGGTACCCTCGAG TC | Amplify pACYC-Duet backbone for cloning of *drmA* insert | fw |
| BN1064 | TATATATACATATGATGGATGAACTC TTAGATGC | Clone *drmC* into plasmid pET-Duet using RE cloning-NdeI | fw |
| BN1065 | ATATATATCTCGAGTTAAATCAGACT AATGATATTTGTG | Clone *drmC* into plasmid pET-Duet using RE cloning-XhoI | rv |
| BN172 | AGATCTGCCATATGTATATCTCCTTC | Amplify pET-Duet backbone for cloning of *drmC* insert | rv |
| BN2290 | GATCGCTGACGTCGGTACCCTCGAG TC | Amplify pET-Duet backbone for cloning of *drmC* insert | fw |
| BN3096 | CATACCCTATCTCATTTGTTGAT | Create mutant DrmB-ΔDUF1998 in 13ss and in pCDF | fw |
| BN3095 | TTATCATTACAGCATGGCATATCGGA TAC | Create mutant DrmB-ΔDUF1998 in 13ss and in pCDF | rv |
| BN3104 | TTATCATTATCGCATTGACATAGGTA CAG | Create mutant *drmA*-ΔPA in pCDF | rv |
| BN3105 | GAAGTTGAGTCTGGTGTAC | Create mutant *drmA*-ΔPA in pCDF | fw |
| BN3436 | TTGGTGCCCAGTAGCCTC | Create mutant *drmA*-Δloop in pACYC and pCDF | rv |
| BN3437 | TTCGGCTGCTTCACGTGG | Create mutant *drmA*-Δloop in pACYC and pCDF | fw |
| BN3377 | CAACACGACTGGTAGCCTGAATATAC | Amplify DNA from pCDF-Duet with *drmA*-ΔPA and clone into pACYC-Duet | rv |
| BN3378 | GGACACCAAATTTATCAAGGACGATC | Amplify DNA from pCDF-Duet with *drmA*-ΔPA and clone into pACYC-Duet | fw |
| BN3375 | GTATATTCAGGCTACCAGTCGTGTTG | Amplify DNA from synthetic block to clone in 13SS and pCDF-Duet for *drmA* mutants R1294A, V1296G, V1296W | fw |
| BN3376 | GATCGTCCTTGATAAATTTGGTGTCC | Amplify DNA from synthetic block to clone in 13SS and pCDF-Duet for *drmA* mutants R1294A, V1296G, V1296W | rv |
| BN3377 | CAACACGACTGGTAGCCTGAATATAC | Amplify vectors 13SS and pCDF-Duet DNA to insert amplified DNA of *drmA* mutants R1294A, V1296G, V1296W | rv |
| BN3378 | GGACACCAAATTTATCAAGGACGATC | Amplify vectors 13SS and pCDF-Duet DNA to insert amplified DNA *drmA* mutants R1294A, V1296G, V1296W | fw |

| | | | |
|---|---|---|---|
| BN3379 | GATGAACAGAACTTGAATGTGAGC | Amplify DNA from synthetic block to clone in 13SS and pCDF-Duet for *drmA* mutants R569D, Q572D, K610D, R658D, R649A, K803A, R810A | rv |
| BN3380 | GATAATTAAATCAGGCGGTCGGATA G | Amplify DNA from synthetic block to clone in 13SS and pCDF-Duet for *drmA* mutants R569D, Q572D, K610D, R658D, R649A, K803A, R810A | fw |
| BN3381 | GCTCACATTCAAGTTCTGTTCATC | Amplify vectors 13SS and pCDF-Duet DNA to insert amplified DNA *drmA* mutants R569D, Q572D, K610D, R658D, R649A, K803A, R810A | rv |
| BN3382 | CTATCCGACCGCCTGATTTAATTATC | Amplify vectors 13SS and pCDF-Duet DNA to insert amplified DNA *drmA* mutants R569D, Q572D, K610D, R658D, R649A, K803A, R810A | fw |

### Supplementary Table 4 | Plasmids used in this study.

| Plasmid | Description | Name in paper | Antibiotic resistance marker | Source |
|---|---|---|---|---|
| pTU515 | 13SS-*drmB* fused to His-tag | | Spectinomycin | This paper |
| pTU518 | 13SS-*drmB*-ΔDUF1998 fused to His-tag | | Spectinomycin | This paper |
| pTU516 | pACYC-Duet-*drmA* | | Chloramphenicol | This paper |
| pTU520 | pACYC-Duet-*drmA*(ΔPA) | | Chloramphenicol | This paper |
| pTU542 | pACYC-Duet-*drmA*(Δloop) | | Chloramphenicol | This paper |
| pTU522 | pACYC-Duet-*drmA*(V1296G) | | Chloramphenicol | This paper |
| pTU523 | pACYC-Duet-*drmA*(V1296W) | | Chloramphenicol | This paper |
| pTU524 | pACYC-Duet-*drmA*(R569D) | | Chloramphenicol | This paper |
| pTU525 | pACYC-Duet-*drmA*(Q572D) | | Chloramphenicol | This paper |
| pTU526 | pACYC-Duet-*drmA*(K610D) | | Chloramphenicol | This paper |
| pTU527 | pACYC-Duet-*drmA(*R658D) | | Chloramphenicol | This paper |
| pTU528 | pACYC-Duet-*drmA*(R649A) | | Chloramphenicol | This paper |
| pTU529 | pACYC-Duet-*drmA*(K803A) | | Chloramphenicol | This paper |
| pTU530 | pACYC-Duet-*drmA*(R810A) | | Chloramphenicol | This paper |
| pTU531 | pACYC-Duet-*drmA*(R1294A) | | Chloramphenicol | This paper |
| pTU517 | pET-Duet-*drmC* | | Ampicillin | Aparicio-Maldonado et al. |
| pTU495 | pCDF-Duet-*drmABC* | | Streptomycin | Aparicio-Maldonado et al. |
| pTU519 | pCDF-Duet-*drmABC* with *drmB*-ΔDUF1998 | | Streptomycin | This paper |
| pTU521 | pCDF-Duet-drmABC with *drmA*(ΔPA) | | Streptomycin | This paper |
| pTU543 | pCDF-Duet-drmABC with *drmA*(Δloop) | | Streptomycin | This paper |
| pTU532 | pCDF-Duet-drmABC with *drmA*(V1296G) | | Streptomycin | This paper |
| pTU533 | pCDF-Duet-drmABC with *drmA*(V1296W) | | Streptomycin | This paper |
| pTU534 | pCDF-Duet-drmABC with *drmA*(R569D) | | Streptomycin | This paper |

| | | | | |
|---|---|---|---|---|
| pTU535 | pCDF-Duet-drmABC with *drmA*(Q572D) | | Streptomycin | This paper |
| pTU536 | pCDF-Duet-drmABC with *drmA*(K610D) | | Streptomycin | This paper |
| pTU537 | pCDF-Duet-drmABC with *drmA(*R658D) | | Streptomycin | This paper |
| pTU538 | pCDF-Duet-drmABC with *drmA*(R649A) | | Streptomycin | This paper |
| pTU539 | pCDF-Duet-drmABC with *drmA*(K803A) | | Streptomycin | This paper |
| pTU540 | pCDF-Duet-drmABC with *drmA*(R810A) | | Streptomycin | This paper |
| pTU541 | pCDF-Duet-drmABC with *drmA*(R1294A) | | Streptomycin | This paper |

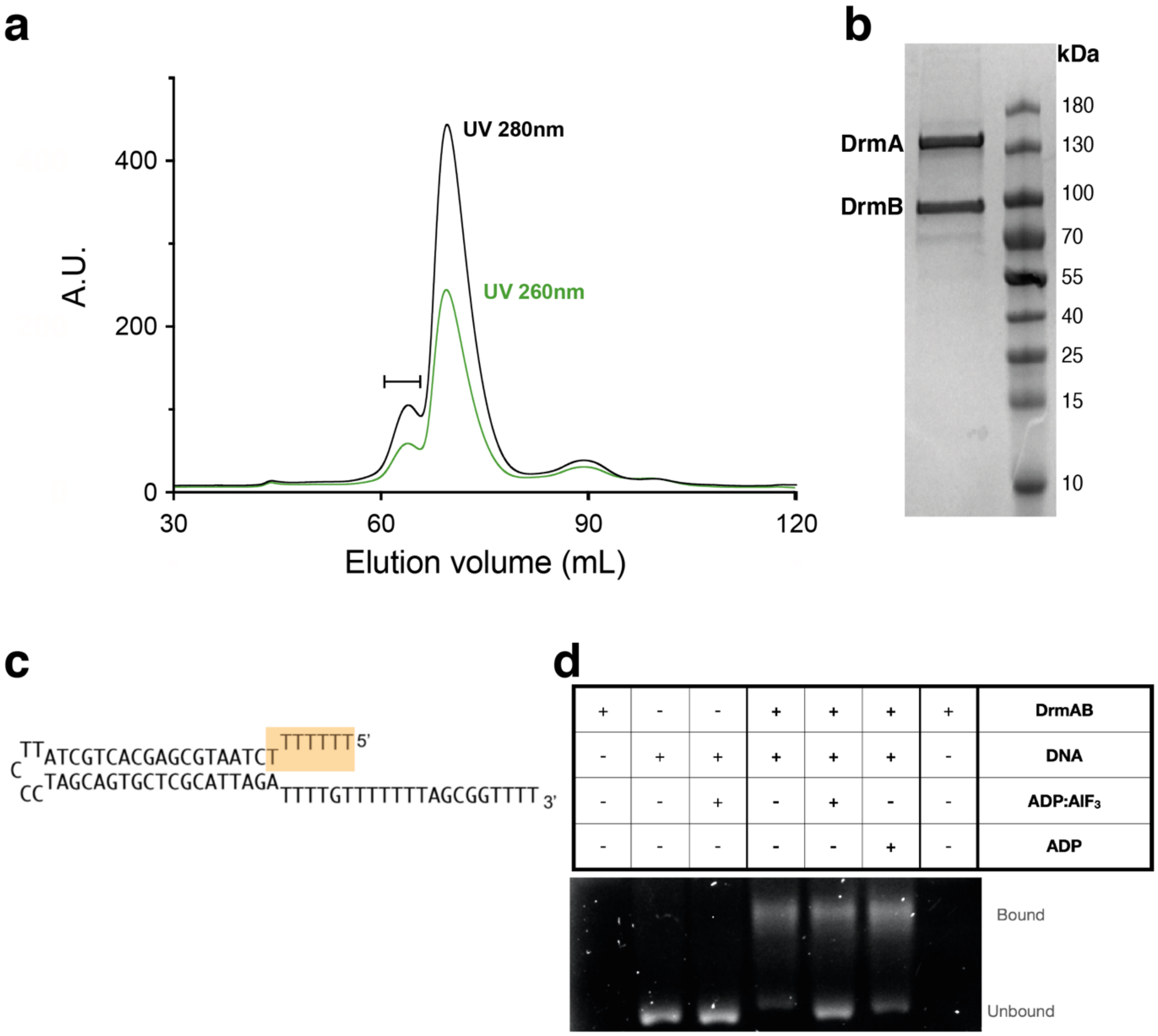


**Supplementary Fig. 1 I DNA binding by DrmAB**

**a**, Size-exclusion chromatogram of DrmAB complex. DrmB was tagged and used to pull down co-expressed DrmAB complex. Line denotes DrmAB species analyzed in panel B. **b**, SDS PAGE analysis of DrmAB complex purified by size-exclusion chromatography. DrmAB assembles with a 1:1 DrmA:DrmB stoichiometry. **c**, Secondary structure diagram of DNA stem-loop used for structural analyses (**Table 2**). 5' region visible in cryo-EM reconstruction is denoted by orange box. **d**, Native gel shift assay used to determine sample preparation conditions for cryo-EM. 4 $\mu$M DNA hairpin (**Table 2**) was heat annealed and incubated with 10 $\mu$M DrmAB in the absence or presence of 1 mM ADP or ADP:$AlF_3$. Bound and free DNA were separated using EMSA.

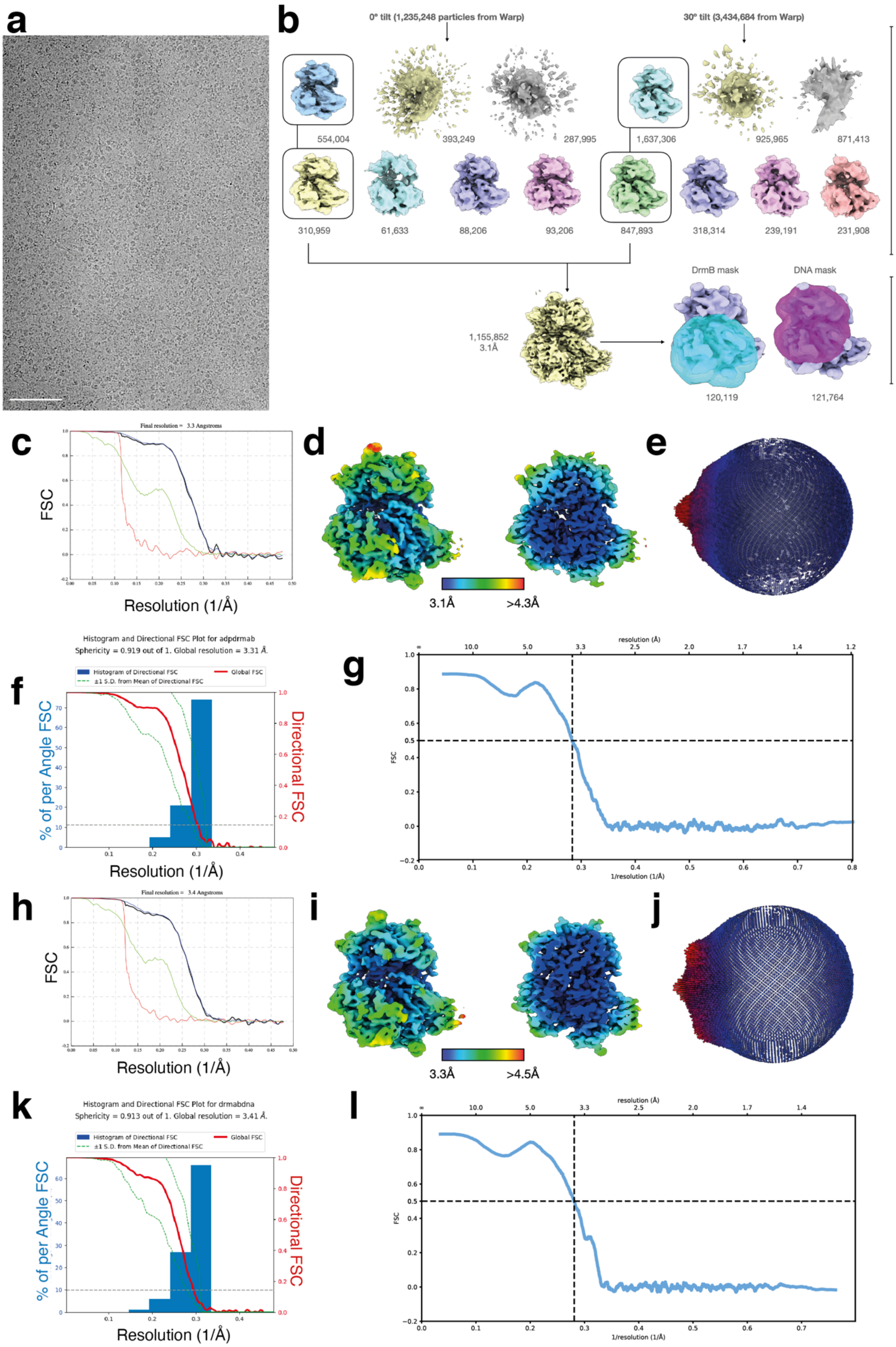

a
b
0° tilt (1,235,248 particles from Warp)
30° tilt (3,434,684 from Warp)
554,004
393,249
287,995
1,637,306
925,965
871,413
310,959
61,633
88,206
93,206
847,893
318,314
239,191
231,908
CryoSparc
DrmB mask
DNA mask
1,155,852
3.1Å
120,119
121,764
Relion 3.1
c
Final resolution = 3.3 Angstroms
FSC
Resolution (1/Å)
d
3.1Å
>4.3Å
e
f
Histogram and Directional FSC Plot for adpdrmab
Sphericity = 0.919 out of 1. Global resolution = 3.31 Å.
Histogram of Directional FSC
±1 S.D. from Mean of Directional FSC
Global FSC
% of per Angle FSC
Directional FSC
Resolution (1/Å)
g
resolution (Å)
FSC
1/resolution (1/Å)
h
Final resolution = 3.4 Angstroms
FSC
Resolution (1/Å)
i
3.3Å
>4.5Å
j
k
Histogram and Directional FSC Plot for drmabdna
Sphericity = 0.913 out of 1. Global resolution = 3.41 Å.
Histogram of Directional FSC
±1 S.D. from Mean of Directional FSC
Global FSC
% of per Angle FSC
Directional FSC
Resolution (1/Å)
l
resolution (Å)
FSC
1/resolution (1/Å)

**Supplementary Fig. 2 | Cryo-EM data processing of DrmAB-ADP-DNA dataset**

**a,** Representative cryo-electron micrographs of DrmAB:ADP:DNA. Scale bar – 100 nm. **b,** Data processing workflow detailing how tilted and untilted particle stacks were combined to a consensus reconstruction. While further classification of said reconstruction yielded a 2.8Å-resolution map, the quality of density corresponding to both ssDNA and DrmB N-terminal half were poor. To ameliorate this, focused 3D classification using either a DrmB mask or a DNA mask was used. **c,** Gold-standard Fourier shell correlation (FSC) curve for DrmAB:ADP. **d,** EM density map of DrmAB:ADP color-coded according to local resolution. **e**, Angular distribution plot calculated in Relion of DrmAB:ADP. **f**, Directional 3D FSC for DrmAB:ADP calculated by 3DFSC[44]. **g**, Map-to-model FSC for DrmAB:ADP. **h,** Gold-standard Fourier shell correlation (FSC) curve for DrmAB:ADP:DNA. **I:** EM density map of DrmAB:ADP:DNA color-coded according to local resolution. **j**, Angular distribution plot calculated in Relion of DrmAB:ADP:DNA. **K**: Directional 3D FSC for DrmAB:ADP:DNA calculated by 3DFSC[44]. **l**, Map-to-model FSC for DrmAB:ADP:DNA.

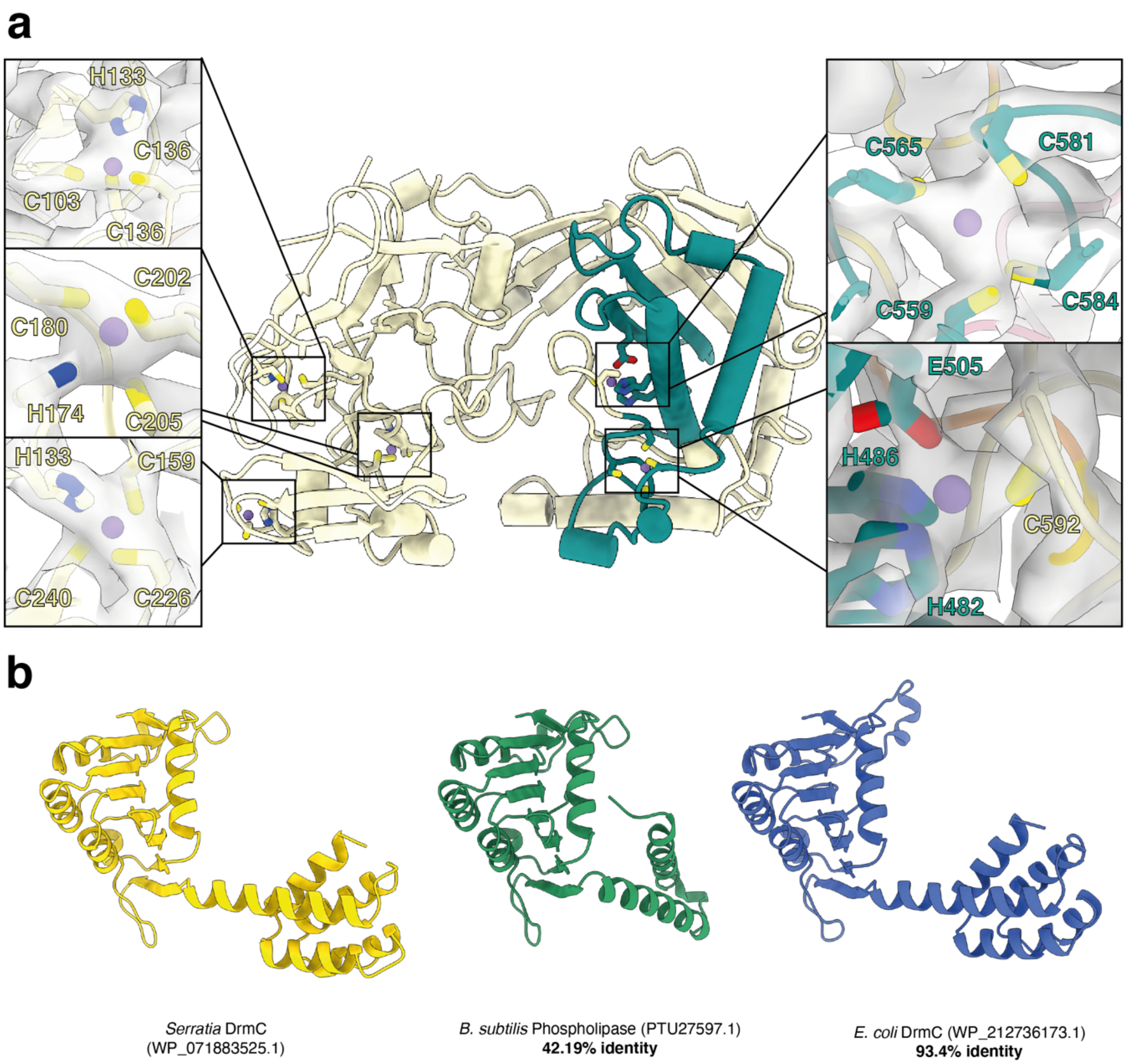


**Supplementary Fig. 3 | Structural analysis of DrmB and DrmC**

**a,** Multiple putative metal coordination sites within DrmB, with representative cryo-EM densities. **b,** Structural models of DrmC from *Serratia* (yellow), a *B. subtilis* PLD-nuclease protein homologue of DrmC (green) and an *E. coli* PLD-nuclease protein homologue of DrmC (blue). The DISARM operon in this study originated in *Serratia*, and *in vivo* assays were performed in *E. coli. B. subtilis* was the model organism used for *in vivo* assays in a previous DISARM study[7]. Structural models were generated using AlphaFold2[45].

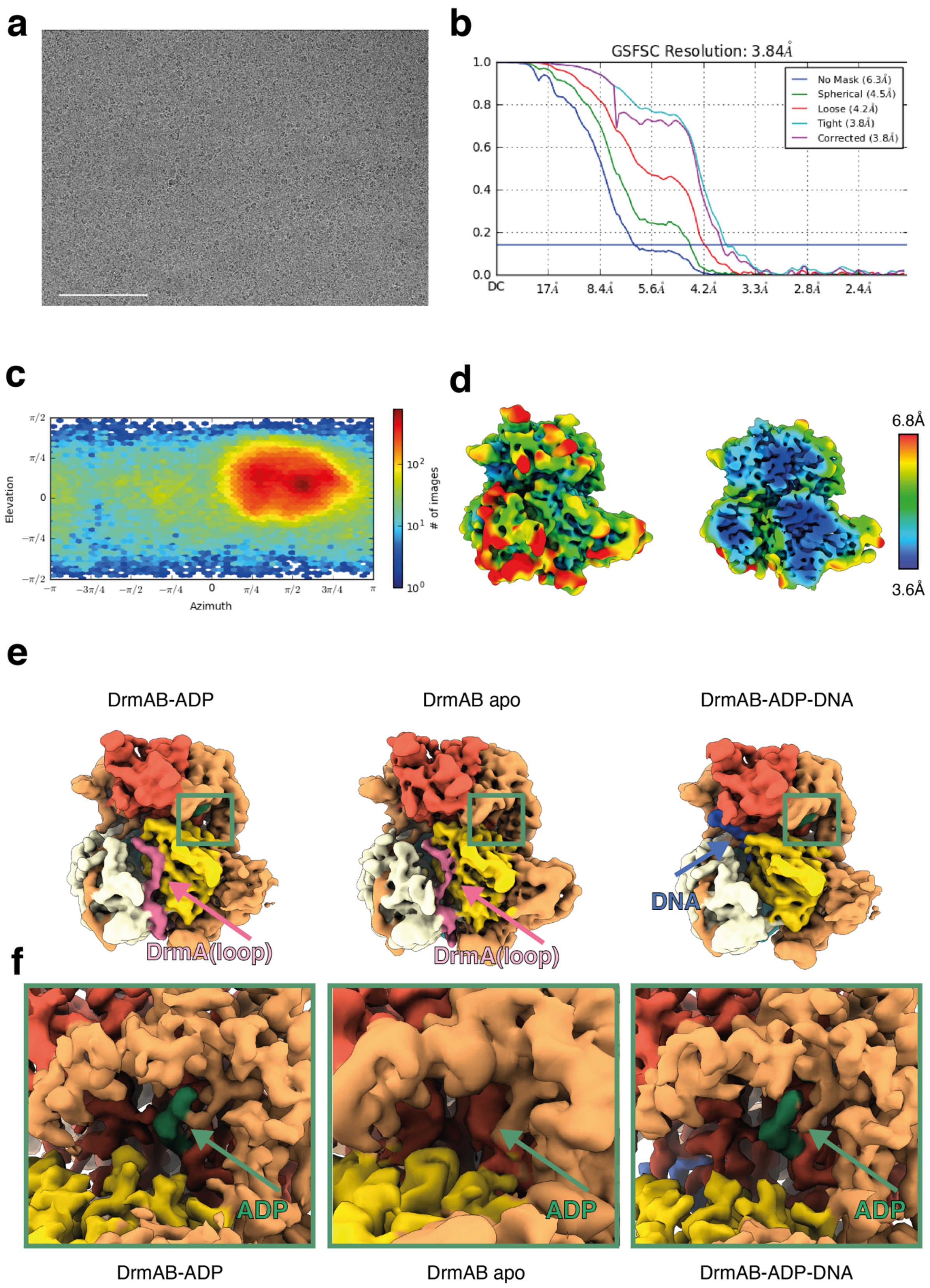


**Supplementary Fig. 4 l Structural features of DrmAB complex and TL**

**a,** Representative cryo-electron micrograph of DrmAB. Scale bar – 100 nm. **b**, Gold-standard FSC curves for DrmAB. **c**, Angular distribution plot for DrmAB calculated in cryoSPARC. **d,** EM density map of DrmAB color-coded according to local resolution. **e**, Presence of TL EM density features within DrmAB:ADP, DrmAB, and DrmAB:ADP:DNA maps. TL is pink, and DNA is blue. Green box corresponds to ATP binding site. **f,** ATP-binding site within DrmA RecA domains. DrmAB apo complex contains no nucleotide density, but still contains TL density.
